# Evaluation of the NCCN guidelines using the RIGHT Statement and AGREE II instrument: a cross-sectional review

**DOI:** 10.1101/442285

**Authors:** Cole Wayant, Craig Cooper, D’Arcy Turner, Matt Vassar

## Abstract

**Introduction:** Robust, clearly reported clinical practice guidelines (CPGs) are essential for evidence-based clinical practice. The Reporting Items for practice Guidelines in HealThcare (RIGHT) statement and Appraisal of Guidelines for Research and Evaluation (AGREE) II instrument were published to improve the methodological and reporting quality in healthcare CPGs.

**Methods:** We applied the RIGHT statement checklist and AGREE II instrument to 48 National Comprehensive Cancer Network (NCCN) guidelines. Our primary objective was to assess the adherence to RIGHT and AGREE II items. Since neither RIGHT nor AGREE-II can judge the clinical usefulness of a guideline, our study is designed to only focus on the methodological and reporting quality of each guideline.

**Results:** The NCCN guidelines demonstrated notable strengths and weaknesses. For example, RIGHT statement items 19 (conflicts of interest), 7b (description of subgroups), and 13a (clear, precise recommendations) were fully reported in all guidelines. However, the guidelines inconsistently incorporated patient values and preferences and cost, nor did they consistently describe the method for assessing the quality and certainty of evidence. Regarding the AGREE II instrument, the NCCN guidelines scored highly on the domains 4 (clear, precise recommendations) and 6 (handling of conflicts of interest), but lowest on domain 2 (inclusion of all relevant stakeholders).

**Conclusions:** In this investigation we found that NCCN CPGs demonstrate key strengths and weaknesses with respect to the reporting of key items essential to CPGs. We recommend the continued use of NCCN guidelines and adherence to the RIGHT and AGREE II items. Doing so serves to improve the evidence delivered to healthcare providers, thus potentially improving patient care.

## Introduction

Robust, clearly reported clinical practice guidelines (CPGs) are essential for evidence-based clinical practice. The Institute of Medicine recognizes CPGs as necessary reference material for physicians seeking to optimize patient care.^1^ CPGs are capable of increasing the quality of patient care and improving patient outcomes^2^, but the adoption of low-quality guidelines may result in widespread use of ineffective treatments, inefficient practices, and harm to patients^3,4^. Even though they are an essential resource, CPGs have historically exhibited low-quality reporting.^5^ The ramifications of low reporting quality in CPGs are broad, but most pressing is the lack of a distinction between poor methods and poorly reported methods. In practice, the two may be indistinguishable. For example, if CPG developers perform a narrow, inadequate search of the literature, their subsequent recommendations may not be reproducible or trustworthy. Similarly, if the CPG developers do not report their search strategy, the question remains as to whether the recommendations are trustworthy. The quality of CPG reporting is as important as its methodological quality.

In oncology, new drug approvals may result in rapid changes to patient care. Articulating the available evidence, its strength, and its limitations to physicians is vital. The National Comprehensive Cancer Network (NCCN) — arguably the premier guideline organization in the United States^6^ — has a policy to update their CPGs “at least annually.”^7^ This policy of annual updates highlights the urgent need for clear reporting of current and future CPGs.

Two popular instruments exist for assessing the quality of CPGs in healthcare: The Reporting Items for practice Guidelines in HealThcare (RIGHT) statement^8^ and the Appraisal of Guidelines for Research and Evaluation (AGREE) II instrument^9^. The AGREE-II instrument includes items related to the methodological (e.g., quality of search strategy, inclusion of stakeholder preferences) and reporting quality of CPGs, whereas the RIGHT statement focuses solely on reporting quality (e.g., providing a summary of recommendations, disclosure of funding source). Neither was created as a handbook for developing guidelines. According to the RIGHT Statement authors, the RIGHT Statement is not designed to assess the inherent quality of a guideline.^8^ Rather, the RIGHT Statement is designed to complement tools that are designed to assess the inherent quality of a guideline, such as the AGREE-II instrument.

Given the comprehensiveness and importance of the NCCN CPGs to oncology practice^6^, the aim of this investigation is to highlight the strengths and weaknesses in the reporting of NCCN guidelines. By doing so, we aim to improve the delivery of oncology evidence to oncologists and improve patient care. In this study we applied the RIGHT Statement and AGREE-II instrument to 49 NCCN guidelines for the treatment of cancer by cancer site.

## Methods

We retrieved all 49 NCCN guidelines on March 21, 2018. Prior to data extraction, CW, CC, and DT reviewed the RIGHT statement and AGREE-II instrument manuals to become familiar with the checklist items.^8,9^ We met and devised a Google Form for both tools. CW, CC, and DT extracted data for all items from each tool. Since the NCCN does not detail their full methods in each CPG, and provides a full explanation of many aspects of their methods on their website (www.nccn.org), we extracted data from the CPG and website policy documents. Any discrepancies in data extraction were resolved via consensus discussion. After extraction and validation of all Google Form responses, we exported these responses to a Google Sheet. We used this Google Sheet to calculate summary statistics.

The design of the RIGHT Statement parallels other statements and reporting guidelines, such as CONSORT for clinical trials or PRISMA for systematic reviews, and consists of a 35-item checklist and an Explanation and Elaboration document.^8^. For each of the items we assigned a numeric score of 1 (full adherence), 0.5 (partial adherence), or 0 (no adherence). An example of partial adherence may be if a a guideline provides a partial explanation of cancer epidemiology. Full explanation includes a description of prevalence/incidence, morbidity, mortality, and burden (including financial). We present summary data using the described scoring convention for each of the 35-items.

The AGREE-II instrument is organized differently, and consists of 23 items divided into 6 domains, with each item scored on a 1 (strongly disagree) to 7 (strongly agree) Likert-type scale. A description of each domain is available in Table 1. In accordance with the AGREE-II manual,^9^ we calculated a scaled domain score for each domain for each CPG. The scaled domain score is calculated as follows:

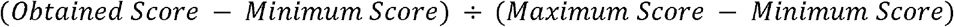

The scaled domain score can be converted to an average rating (1 to 7 scale) by multiplying the scaled domain score by 7. The obtained score is calculated for each domain and is the sum of all rater scores in that domain. The minimum score is calculated by multiplying the minimum item score (1, strongly disagree), the number of raters (3, in this study), and the number of items in the domain. The maximum score is calculated similarly, but substitutes the maximum item score (7, strongly agree) for the minimum item score. Lastly, we made a judgement about whether the CPG should be recommended or not based largely on the 6 scaled domain scores for each CPG.

**Table 1.**
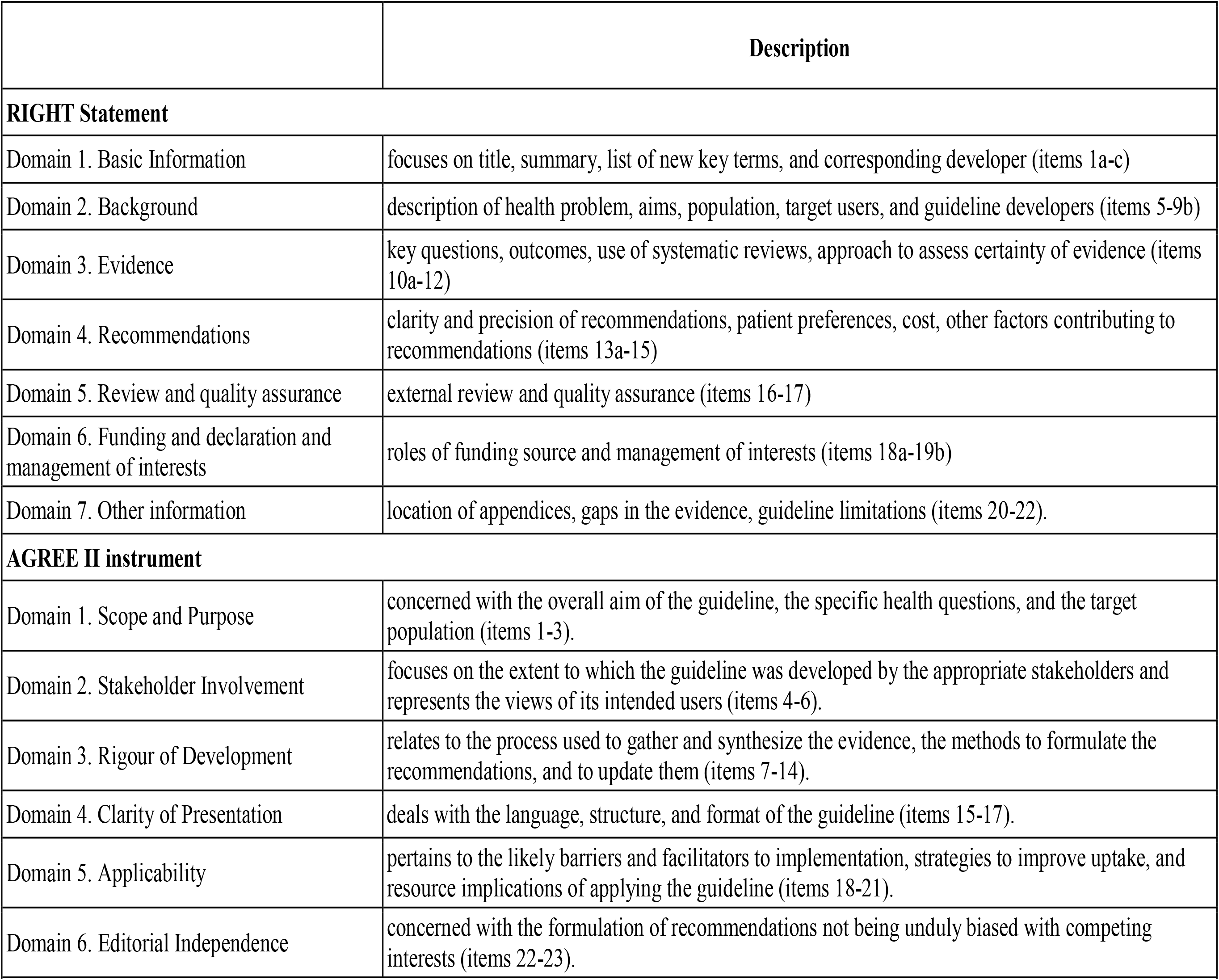
Description of the seven RIGHT statement and six AGREE II instrument domains.

Our primary objective was to assess CPG scores on the RIGHT statement and AGREE-II instrument. Since all NCCN guidelines were published after the RIGHT statement and AGREE-II instrument were published, they are all eligible for analysis. As neither the RIGHT statement or AGREE-II instrument can judge the clinical usefulness of a guideline, our study is designed to only focus on the methodological and reporting quality of each guideline.

## Results

We identified 49 NCCN CPGs for the treatment of cancer by site. The Uveal Melanoma CPG was excluded because the Discussion section (the narrative section of NCCN guidelines) is under development and not written.

### RIGHT Statement

The NCCN guidelines were largely homogenous, and many key methodological items were reported clearly in policy documents on the NCCN website. The standard deviation for 30 of the 35 items (85.7%) was zero. Table 2 shows each NCCN guideline and its average score for all RIGHT statement items.

**Table 2.**
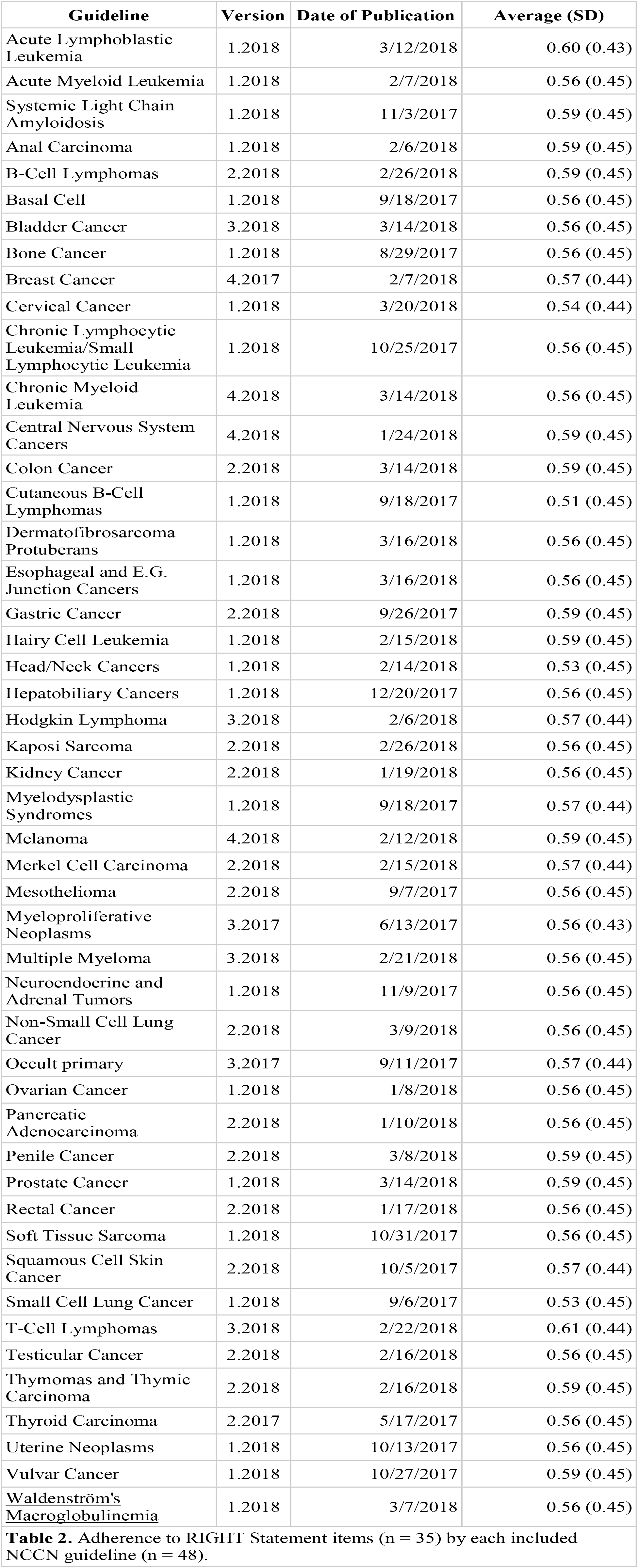
Adherence to RIGHT Statement items (n = 35) by each included NCCN guideline (n = 48).

Notable strengths of the NCCN CPGs were the reporting of conflicts of interest for all authors (items 19a and 19b), complete description of pertinent subgroups (item 7b), and the clarity of CPG recommendations (item 13a). Notable deficiencies were the description of stakeholder involvement (e.g., patient views and preferences) [item 14a], the cost and resource implications of therapies (item 14b), which outcomes were prioritized when formulating recommendations (item 10b), and the approach to assess the certainty of the quality of evidence (item 12).

### AGREE-II

Table 3 shows the scaled domain scores for each NCCN CPG. Using the AGREE-II instrument we were able to assess CPG scores in six domains, each essential to a methodologically robust CPG. No guideline scored extremely low for any domain. The fourth domain (Clarity of Presentation) and sixth domain (Editorial Independence) scored the highest, overall. The Clarity of Presentation domain asks whether the recommendations are specific and unambiguous, if alternative treatment options were mentioned, and if the key recommendations are easily identifiable. The sixth domain asks questions about the influence of the funding source on CPG development and whether conflicts of interest were disclosed. The lowest, individual domain score was 36.1% in the Applicability domain for the acute lymphoblastic leukemia CPG. This score indicates that average score (1 to 7 scale) for this domain was approximately 2.5. With respect to overall domain scores across all guidelines, the Stakeholder Involvement domain scored the lowest with an average score of 48.6% (e.g., 3.4 out of 7). The Stakeholder Involvement domain asks questions related to the description of guideline development members, the incorporation of target population views and preferences, and the identification of target users of the guidelines.

**Table 3.**
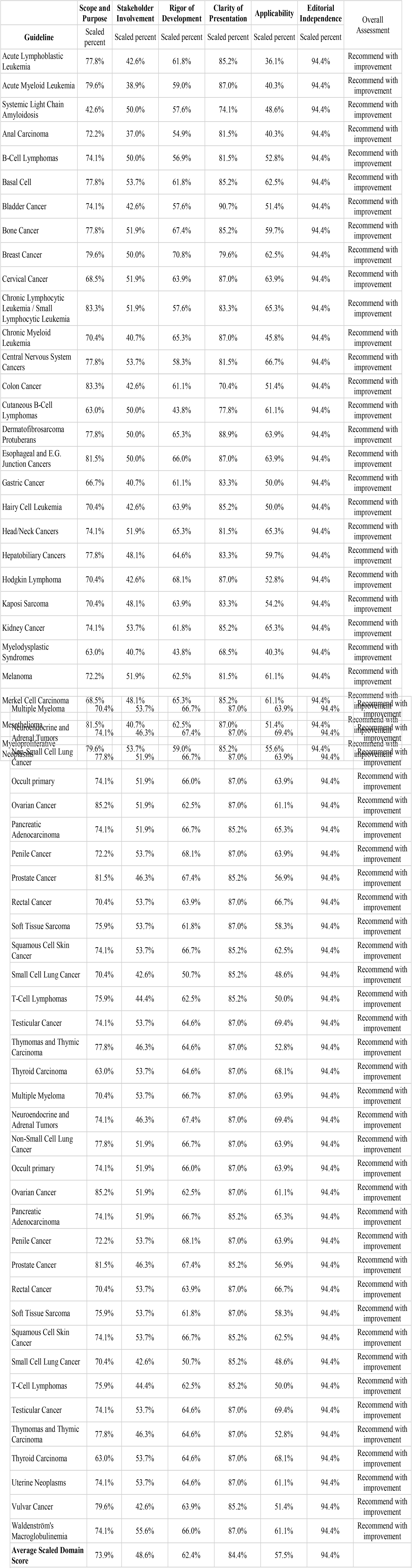
Scaled AGREE II domain score for all included guidelines (n = 48).

## Discussion

In this investigation we found that NCCN CPGs demonstrate key strengths and weaknesses with respect to the reporting of key items essential to CPGs. For example, the NCCN CPGs require conflicts of interest disclosure, clearly describe all pertinent subgroups, and delineate key recommendations. On the other hand, the NCCN CPGs did not consistently describe how patient values and preferences were incorporated into recommendations, the financial burden of the recommendations, or describe the approach used to assess the certainty of the evidence underpinning the recommendations.

The AGREE-II instrument^9^ was developed to assess CPG quality in six, equally essential domains ranging from describing the purpose of the CPG to the applicability of the CPG recommendations. We found that they scored well enough to continue being recommended in clinical practice, but key methodological items were not reported, thus highlighting areas where the delivery of oncology evidence can be improved. Through applying the RIGHT Statement, which was created to be used alongside the AGREE-II instrument, we confirmed that improvements in the reporting of several key items would strengthen the impact of NCCN CPGs by increasing the clarity and comprehensiveness of the recommendations.

None of the NCCN CPGs described the process by which patient values and preferences were solicited and incorporated into the guideline recommendations. The primary reason for incorporating patient values and preferences into CPG recommendations is that recommendations that are aligned with patient values may be more easily adopted and implemented^10–12^. Until recently, there were no firmly established processes for including patient values and preferences in CPG recommendations. To address this gap, the GRADE (Grading of Recommendations Assessment, Development and Evaluation) working group created the GRADE Evidence-to-Decision (EtD) framework^11^. Previously, the GRADE approach has been used to assess the quality and certainty of evidence underpinning clinical practice guideline recommendations. The NCCN CPGs do not currently use the GRADE approach, rather they seem to rely on guideline development member assessments of the quality of evidence. Adoption of the GRADE approach would improve the objectivity, applicability, and comparability of NCCN recommendations. Concurrent adoption of the GRADE EtD framework would ensure the incorporation of patient values and preferences in all recommendations.

Additional, minor adjustments to the reporting of NCCN CPGs would improve the delivery of oncology evidence. First, stating key research questions that formed the basis for treatment recommendations in PICO (Population, Intervention, Comparator, Outcome) format would guide physicians through the purpose and scope of the guideline^13–15^. It may be that, due to how comprehensive the NCCN CPGs are, that listing all PICO-format questions is not practical. Should this be the case, we recommend including a section in the CPG that clearly describes the scope, limitations, and gaps in the NCCN recommendations. A second, related adjustment includes listing the outcomes that were most important when developing the CPG recommendations. For example, if efficacy outcomes are the primary basis for the recommendations, or recommending one treatment over another, physicians would benefit from that understanding.

In conclusion, we recommend the continued use of NCCN CPGs to guide oncologists in patient care. We have outlined key recommendations that would improve the completeness of reporting and increase transparency. These recommendations include the adoption of the GRADE and GRADE-EtD approach, describing key questions in PICO format, and sorting which outcomes were important when developing recommendations. We believe that adopting these recommendations will not only improve the NCCN CPGs, but oncology clinical care as well.

